# Cryo-electron tomography reveals structural insights into the membrane binding and remodeling activity of dynamin-like EHDs

**DOI:** 10.1101/2021.09.20.461157

**Authors:** Arthur A. Melo, Thiemo Sprink, Jeffrey K. Noel, Elena Vázquez Sarandeses, Chris van Hoorn, Justus Loerke, Christian M. T. Spahn, Oliver Daumke

**Affiliations:** Max-Delbrück-Centrum for Molecular Medicine, Crystallography, Robert-Rössle-Straße 10, 13125 Berlin, Germany; Freie Universität Berlin, Institute of Chemistry and Biochemistry, Takustraße 6, 14195 Berlin, Germany; Cryo-EM Core Facility, Charité - Universitätsmedizin Berlin at the MDC, Robert-Rössle-Straße 10, 13125 Berlin, Germany; Institut für Medizinische Physik und Biophysik, Charité - Universitätsmedizin Berlin, 10117 Berlin, Germany

**Author notes:** Correspondence and requests for materials should be addressed to A.A.M. or O.D.

**Keywords:** EHD ATPases, dynamin family, oligomerization, membrane remodeling, cryo-electron tomography, subtomogram averaging, membrane binding site

## Abstract

Dynamin-related Eps15-homology domain containing proteins (EHDs) oligomerize on membrane surfaces into filaments leading to membrane remodeling. EHD crystal structures in an open and a closed conformation were previously reported, but structural information on the membrane-bound EHD oligomeric structure has remained enigmatic. Consequently, mechanistic insight into EHD-mediated membrane remodeling is lacking. Here, by using cryo-electron tomography and subtomogram averaging, we determined the structure of an EHD4 filament on a tubular membrane template at an average resolution of 7.6 Å. Assembly of EHD4 is mediated via interfaces in the G-domain and the helical domain. The oligomerized EHD4 structure resembles the closed conformation, where the tips of the helical domains protrude into the membrane. The variation in filament geometry and tube radius suggests the AMPPNP-bound filament has a spontaneous curvature of approximately 1/70 nm^-1^. Combining the available structural and functional data, we propose a model of EHD-mediated membrane remodeling.

## Introduction

Eps15-homology domain containing proteins (EHDs) comprise a eukaryotic family of dynamin-related ATPases (*1*). Mammals contain four closely related EHD homologues (*2*), while only a single member is present in *Drosophila* (termed PAST-1) and *C. elegans* (Rme-1). Rme-1 mediates the release of cargo receptors from the endocytic recycling compartment (*3, 4*). A similar function was demonstrated for mammalian EHD1 and EHD3 (*5, 6*), which also function in the formation of ciliary vesicles (*7, 8*). Furthermore, a conserved tethering complex including EHD1 was shown to coordinate vesicle scission and fusion at sorting endosomes (*9*). EHD2 assemble at the neck of caveolae in ring-like oligomers (*10-15*) that control cellular fatty acid uptake (*12, 16*). EHD4/Pincher mediates macropinocytosis required for retrograde endosomal Trk signaling (*17, 18*). EHD4 also recruits EHD1 to sorting endosomes via hetero-dimerization (*19*). Most recently, a role of an EHD4 complex in the trafficking of vascular endothelial cadherin (VE-cadherin) during angiogenesis was revealed (*20*).

EHDs harbor a dynamin-related GTPase (G) domain that binds to adenine rather than guanine nucleotides (*21, 22*). EHDs tubulate negatively-charged liposomes in an ATP-dependent fashion by the formation of ring-shaped and helical oligomers on the remodeled membranes (*22-24*). Similar to other dynamin superfamily members, oligomerization on membranes stimulates nucleotide hydrolysis (*22, 25*). In reconstitution experiments, ATP hydrolysis in EHD1 induce bulges in tubular membrane templates, leading to membrane scission (*25*).

The crystal structure of the EHD2 dimer in the presence of a non-hydrolyzable ATP analogue revealed a dynamin-related extended G-domain that mediates stable dimerization via an EHD family-specific dimerization interface (interface-1) (*22*). Residues at the N- and C-terminal ends of the G-domain form a composite helical domain. In the crystal structure of the reported EHD2 dimer, the two helical domains protrude in parallel away from the G-domains. This orientation was termed the ‘closed’ conformation of EHDs. The tip of the helical domain was shown to constitute the primary membrane-binding site (*23*).

The C-terminal EH domains interact with linear peptide sequences containing Asn-Pro-Phe (NPF) motifs (*26*) that are present in binding partners, such as MICAL-L1 (*27, 28*), Rabenosyn-5 (*9, 29*), EHBP1 (*30*) and PACSIN1/2 (*13, 15, 31*). In the EHD2 dimeric structure, the EH domains bind back to a Gly-Pro-Phe (GPF) motif in the opposing monomer. In this orientation, the C-terminal tails of the EH domains are positioned into the nucleotide-binding site and block the G-interface. The EH domains may thereby auto-inhibit EHD assembly (*22*).

The crystal structure of an N-terminal deletion variant of EHD4 in the presence of a non-hydrolysable ATP analogue revealed a 50° rotation of the helical domains compared to the EHD2 crystal structure (*32*). In this conformation, the two helical domains in the dimer point away from each other. This arrangement was termed the ‘open’ conformation of EHDs. Spectroscopic experiments indicated that EHD2 binds in the open conformation to flat membrane bilayers (*33*). Furthermore, the EH domains were displaced from the G-domain in the EHD4 structure, suggesting that the open conformation is not compatible with EH-domain mediated auto-inhibition (*32*).

An N-terminal sequence stretch folds back into a conserved hydrophobic pocket of the G-domain in the EHD2 structure (*23*). In the presence of membranes, the N-terminal residues were shown to insert into the lipid bilayer. In turn, a flexible loop at the periphery of the G-domain, the ‘KPF loop’, inserts into the hydrophobic pocket of the G-domain and serves as an oligomerization interface. Deletion of N-terminal residues is therefore expected to stabilize the KPF loop in the G-domain pocket and promote oligomerization. Accordingly, enhanced membrane recruitment and oligomerization was observed for EHD2 and EHD4 variants lacking the N-terminal residues (*23, 32*).

Based on the two available crystal structures, a nucleotide-driven activation model of EHDs has been proposed (*32*). However, since membranes were lacking in any of the reported structures, the detailed conformation of EHDs on membranes and, consequently, their membrane remodeling and oligomerization mode have remained unknown. To address this open issue, we reconstituted an N-terminally truncated EHD4 variant on membrane templates and determined its structure by cryo-electron tomography (cryo-ET) and subtomogram averaging (STA). We thereby clarify the oligomerization mode, reveal how EHD4 interacts with the membrane, demonstrate how the EHD4 oligomer adapts to various membrane curvatures and propose a model for EHD-mediated membrane remodeling.

## Results

### Cryo-ET and subtomogram averaging reveal the structure of the membrane-bound EHD4 oligomer

To understand the molecular mechanisms of EHDs assembly on membranes, EHD4-coated membrane tubules were reconstituted *in vitro*. To this end, we used a previously described N-terminal deletion construct of mouse EHD4 (EHD4^ΔN^, corresponding to amino acids 22-541) that forms a regular protein coat on membranes and displays enhanced membrane recruitment when expressed in eukaryotic cells compared to full length EHD4 (*32*). Full-length EHD4 cannot be expressed in a soluble form in bacteria.

We previously reported that EHD-mediated membrane remodeling is dependent on the presence of ATP (*23, 32*) and, therefore, performed the *in vitro* reconstitutions in the presence of the non-hydrolysable ATP analogue, adenylyl-imidodiphosphate (AMPPNP). Liposomes containing 50% Folch extract from bovine brain, 40% phosphatidylethanolamine and 10% cholesterol were chosen as template since they reproducibly yielded densely coated lipid tubules. These tubules were highly heterogeneous, with luminal diameters ranging between 30 to 100 nm (Fig. 1A and Movie S1) and a variety of different shapes (Fig. S1). The protein coat had a thickness of ∼12 nm (Fig. 1A).

**Figure 1:**
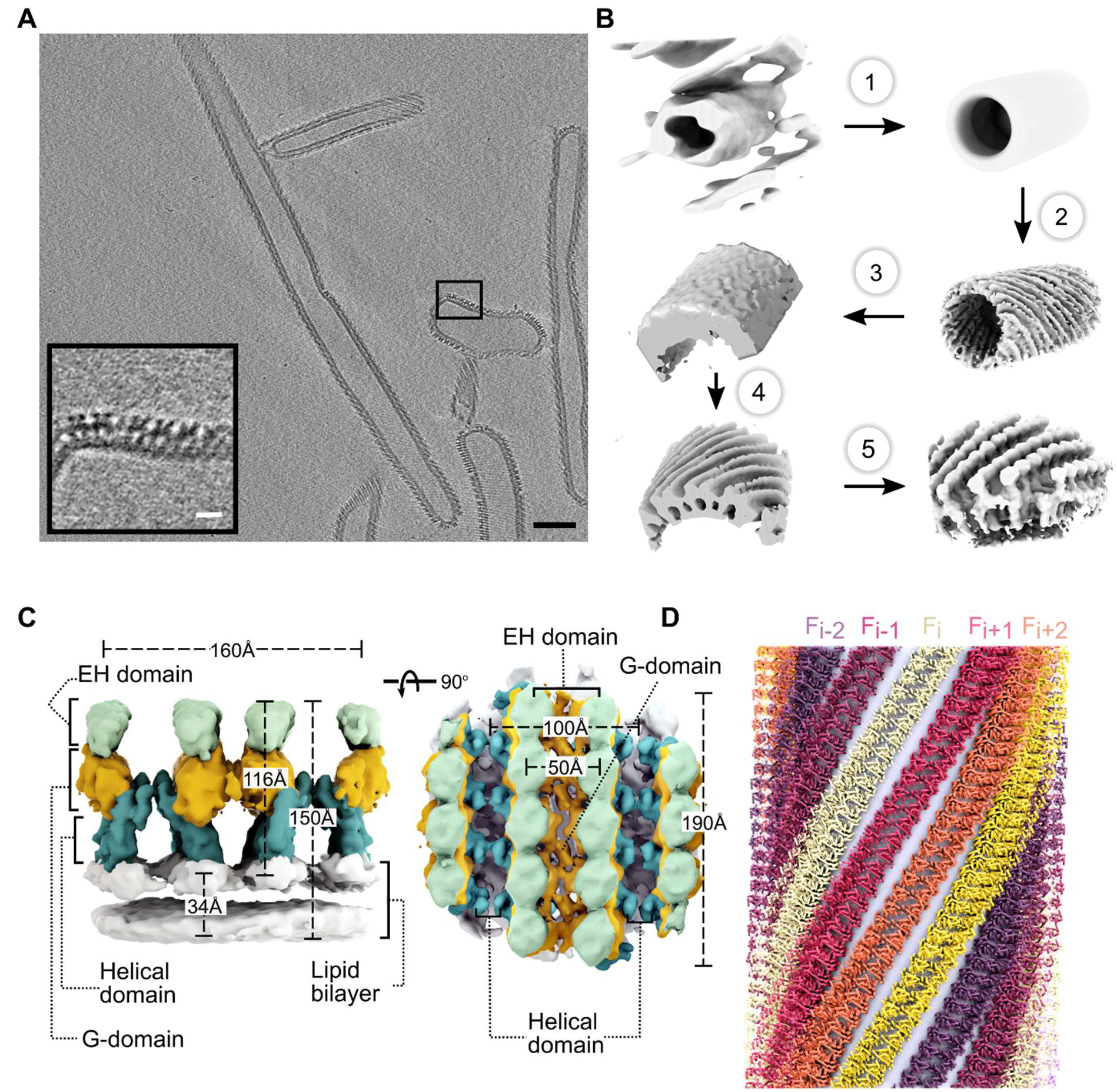
Structure determination of membrane-bound EHD4. **A**. Tomogram reconstruction of EHD4-covered membrane tubules. The boxed area is magnified in the left bottom corner. Scale bars: white 10 nm; black 100 nm. **B**. Subtomogram averaging workflow. Each tubule was individually cropped and averaged (top left). A single tubule shares the same missing wedge with other particles within the same tubule. 1) Random rotation along the particle azimuth to generate the initial template. 2) Subtomogram averaging of individual tubules using Dynamo (see methods for more detail). 3) Subboxing along the tube walls and averaging. 4) Alignment of sub-particles to the template 5) Cropping sub-particles in unbinned tomograms and perform subtomogram averaging. **C**. Subtomogram average of the membrane-bound EHD4^ΔN^ complexed with AMPPNP at 7.6 Å resolution. Domain organization of EHD4 in the filaments in two orientations are shown. EH domains (light green), G-domains (orange), helical domains (teal) and the lipid bilayer (white) are colored individually. **D**. Reconstructed EHD4 right-handed helical filaments wrapping around a membrane tubule. Each filament is indicated in a different color.

To determine the structure of the EHD4 coat, we used cryo-ET and reference-free subtomogram averaging (STA) (Fig. 1B). For this, we collected 56 tilt-series using dose-symmetric tilt scheme, from -60° to 60° with 3° of increment. We divided the data into two half-datasets, which were processed independently, and the resulting structures were compared by Fourier Shell Correlation and averaged together to generate a final structure from 23,813 subtomograms (Table S1, Fig. S2A). The structure was solved at an average resolution of 7.6□Å (Fig. 1C, Fig. S2A, B) and revealed right-handed helical filaments wrapping around lipid tubules (Fig. 1D). The positions of the G-, helical and EH domains could be unambiguously assigned in the density maps by fitting them as rigid bodies, based on the crystal structures (Fig. 1C, Movie S2 and Fig. 2). In the filaments, the EH domains are located furthest from the membrane, the G-domains at the center and the helical domains closest to the membrane (Fig. 1C). An atomic model for the EHD4 filament was obtained using a flexible fitting strategy using the EHD2 crystal structure as a guide (see Methods).

**Figure 2:**
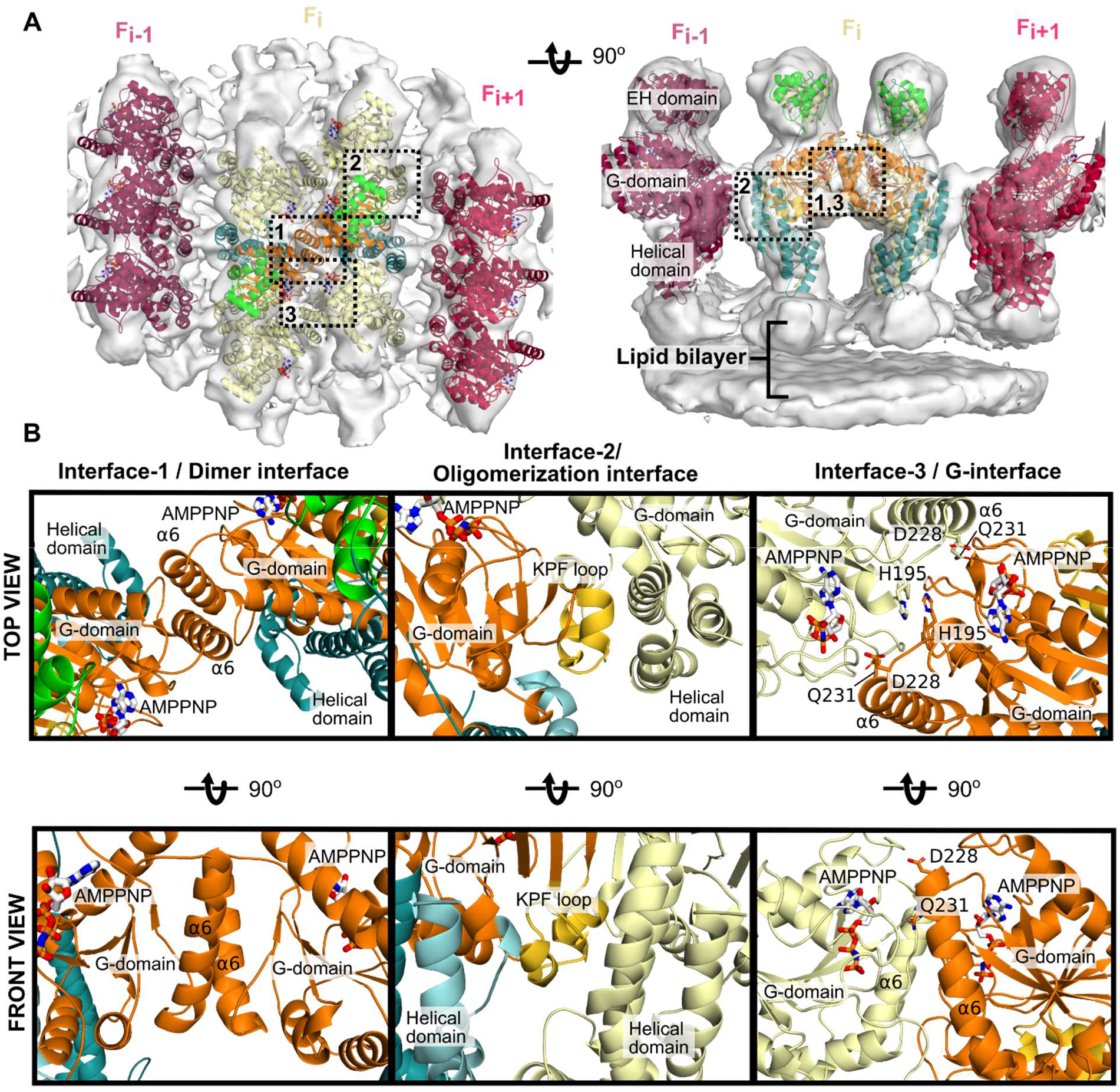
Architecture of the EHD4 filament. **A**. Asymmetric unit of the cryo-ET reconstruction. In the central EHD4 dimer, the domains are colored according to the domain architecture. The cryo-ET density is indicated as grey surface. **B**. The filament is formed by three contacts. Interface-1 comprises a dimer interface in the G-domain, interface-2 is built by a contact between G-domain and helical domain whereas interface-3 represents the G-interface and involves the nucleotide.

### Architecture of the EHD4 filaments

The unit particle used for subtomogram averaging contained three parallel filaments of oligomeric EHD4^ΔN^. The center filament (F_i_) is composed of three EHD4 homodimers, and the adjacent filaments (F_i+1_ and F_i-1_) of three EHD4 monomers each (Fig. 2A). As in the previously reported crystal structures, dimerization of EHD4 is mediated by helix α6 in the G-domain. It forms a highly conserved, two-fold symmetric interface, to which we refer as interface-1 (Fig. 2B).

Each EHD4 dimer interacts in the filament through two additional interfaces (Fig. 2B). Interface-2 is formed between the helical domain of one protomer and the G-domain of the adjacent protomer along the filament. It involves the KPF loop in the G-domain and helices α8 and α12 of the adjacent helical domain. Density of the KPF loop at the periphery of the G-domain dimer is consistent with the open EHD4 crystal structure (Fig. 2A, B). Mutations in this loop were shown to disrupt oligomerization (*32*).

Interface-3 is formed between G-domains of two adjacent dimers across the filament (Fig. 2B) and corresponds to the archetypal G-interface that is conserved in all members of the dynamin family. Dimerization via this interface induces nucleotide hydrolysis in dynamin-related proteins (*34*). Accordingly, mutations in this interface in EHD2 were previously shown to abrogate stimulated ATP hydrolysis (*22*). Highly conserved residues in switch I, switch II, the EHD signature motif (H195) and the N-terminal part of α6 (D228, Q231) are involved in this contact close to the active site.

In the EHD2 crystal structure, the EH domains bind back to the opposing G-domains (*22*). Similarly to this closed conformation, the EH domains were also located on top of the G-domain in our membrane-bound structure (Fig. S3). However, compared to the EHD2 structure, they were shifted away to the periphery (Fig. S3A, C) and stabilized by contacts with EH domains of adjacent dimers (Fig. S3B, D). Accordingly, in this orientation, the C-terminal auto-inhibitory tail may not reach into the active site so that the G-interface can be formed.

### Membrane binding mode of EHD4

We compared the EHD4 monomer to the reported crystal structures of EHD2 and EHD4. Membrane-bound EHD4 adopted the closed conformation of the helical domain, akin to the reported EHD2 conformation (Fig. 3A). In this conformation, the long, central helix α8 from each monomer protrudes towards the membrane.

**Figure 3:**
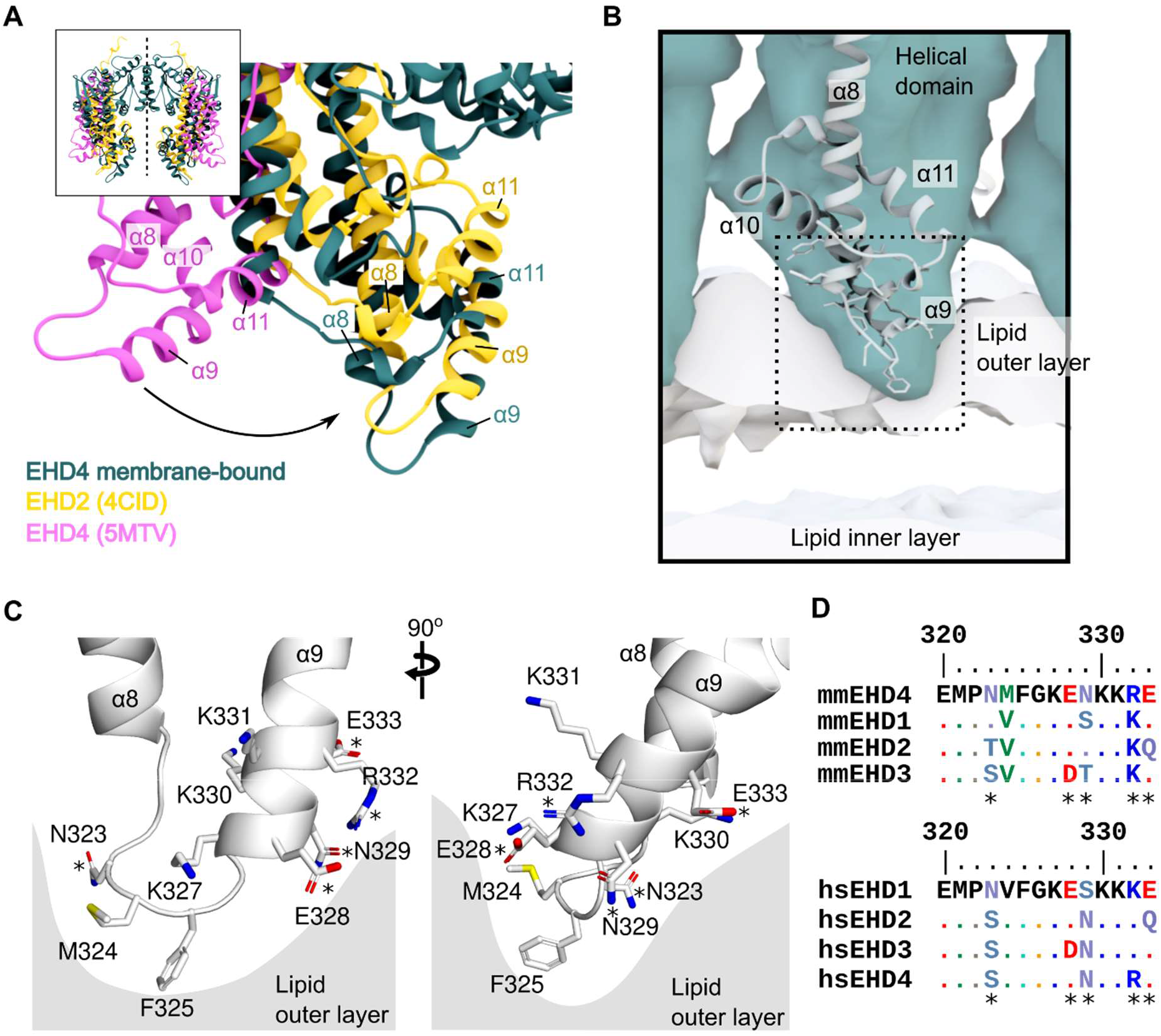
Membrane binding mode of EHD4. **A**. The cryo-ET model of EHD4 (teal) was superimposed with the G-domain on the crystal structures of EHD2 (yellow; pdb 4CID) and EHD4 (pink; pdb 5MTV). The comparison reveals that the membrane-bound structure adopts a closed conformation akin to EHD2. **B**. Membrane binding site of EHD4 insertion into the membrane. **C**. Membrane binding site of EHD4 based on the fittings shown in **A** and **B**. Side chains were modelled based on the EHD4 crystal structure. **D**. Sequence alignment of EHD proteins in mouse (mm - *Mus musculus*) and humans (hs – *Homo sapiens*) reveals a high conservation of the membrane binding site. Conserved residues in all 4 EHD proteins are shown in the sequence alignment are shown as dots (.). Residues that differ in EHDs are highlighted (*).

By electron paramagnetic spin resonance experiments, EHD2 residues at the tip of the helical domain were shown to insert into the membrane (*23*). These findings are consistent with the structure of membrane-bound EHD4. Thus, helices α8 and α9 in the helical domain engage with the membrane bilayer by inserting hydrophobic residues at the connecting loop into the membrane outer leaflet (Fig. 3B, C). In the membrane-bound structure, the Cα atoms of residues K331 and R332 are 1.5-3 Å above the outer leaflet of the bilayer, whereas residues E328, K330 were 0.5-2.5 Å below (Fig. 3C). Residues N323, M324 and F325 at the α8-α9 connecting loop deeply insert into the membrane, with their Cα atom 4, 6, and 8 Å below the membrane density, respectively (Fig. 3C). Charged residues in helix α9, such as K327, K330 and E333 were not inserted into the membrane but were close enough for interaction with the polar lipid head groups (Fig. 3C). Residues in the membrane binding region are highly conserved amongst EHD paralogues and across different species (Fig. 3D). However, each EHD paralogue has a unique combination of membrane-interacting residues (Fig. 3C, D). Thus, EHD proteins bind to membranes through charged residues at helices α8 and α9 and conserved hydrophobic residues at the connecting loop, which likely confer lipid specificity (*22, 35*).

### EHD4 oligomer assembles on membranes of different curvature

EHD4^ΔN^ coated tubules had a wide range of radii (Fig. 4A). Cryo-ET and STA allowed us to probe the architecture of the EHD4^ΔN^ coat on individual tubes, and therefore, allowed the geometry of individual filaments to be discerned. The EHD4^ΔN^ coat was governed by a set of related helical families, which varied in pitch, rise, and subunits per turn (Fig. 4B). By placing the map onto its original position with the refined orientations in the tomograms, we measured a filament’s helical angle as a function of the underlying tubule radius (Fig. 4A, B).

**Figure 4:**
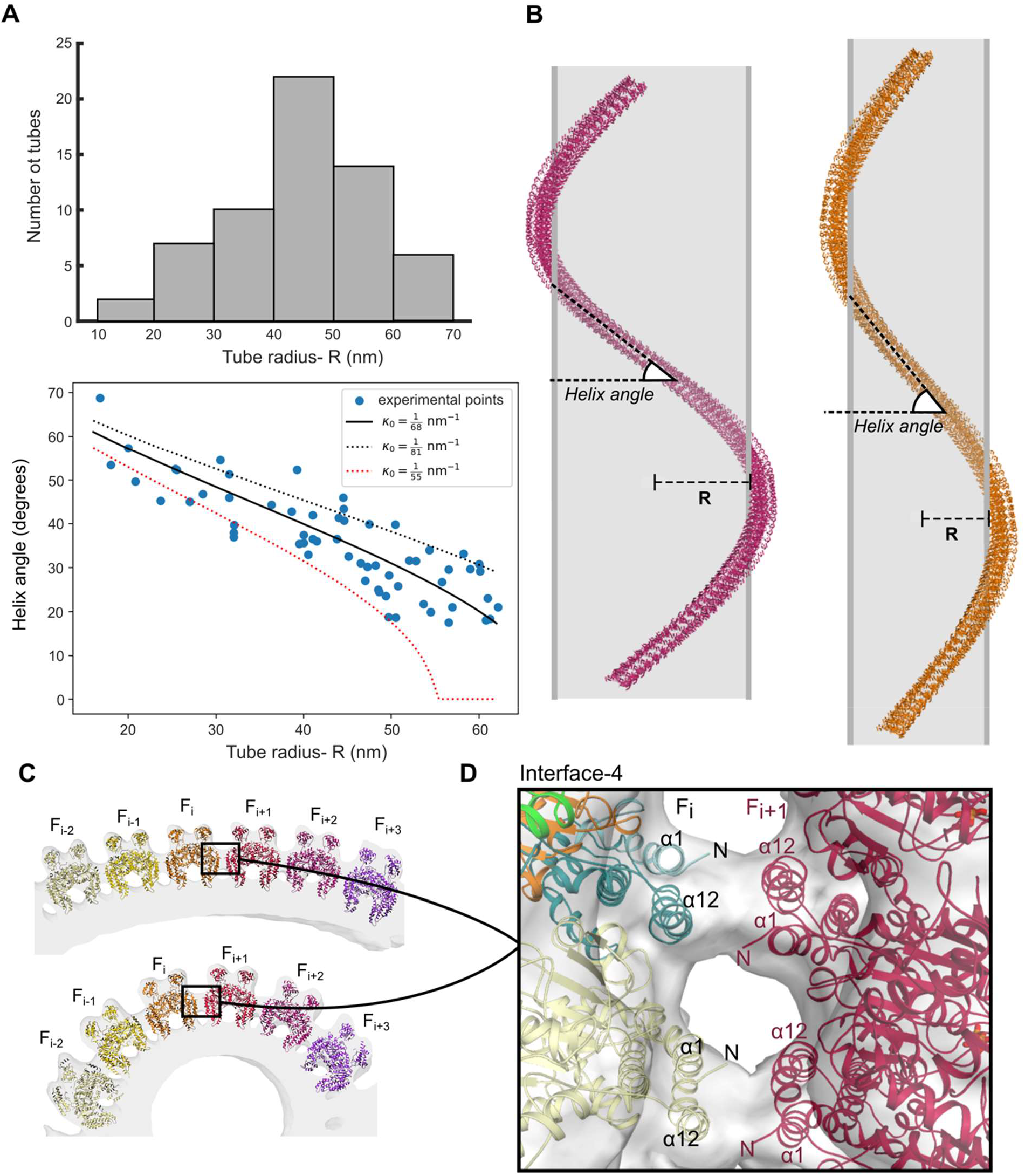
EHD4 filaments adapt to the curvature of membrane tubules. **A**. Distribution of tube radius (top) and the relation of tube diameter and orientation of the filament along the tube axis (bottom). Solid line shows the best fitting spontaneous curvature (see Methods). Dotted lines show the limiting range to illustrate the confidence in predicted **κ**_0_. **B**. EHD4-covered membrane tubes show a wide range of assemblies. A single filament is depicted around the lipid tubule (gray). The average helical assembly of the membrane-bound EHD4 structure is represented on the left and the tube with the smallest diameter is depicted on the right. Both filaments have 42 EHD4 dimers. **C**. EHD4 filaments adopt different conformations in different tubule diameters. The angle between filaments increases in tubules with higher curvature (bottom) along interface-4 (boxed) which acts as a hinge **D**. Interface-4 is formed by contacts between adjacent filaments, which are mediated by helices α1 and α12 of adjacent helical domains. These contacts vary in different cryo-ET reconstructions, indicating a loose interaction that appears to adapt in response to different membrane curvatures.

The helical angle, i.e. the deviation of the helical filament from a simple ring around the tube, tended to increase as the tube radius decreased. This relationship is expected for an elastic filament with a spontaneous curvature less than the curvature of the wrapped tube. Similar behavior is seen for highly constricted dynamin coated tubes (*36, 37*). A best fit suggests that the spontaneous curvature of the AMPPNP-bound-EHD4^ΔN^ filament is 1/68 nm^-1^ (Fig. 4A, bottom). Fitting to more complicated elastic models suggests that the twist stiffness of the filament is significantly weaker than the curvature stiffness and that the angle of each EHD4^ΔN^ dimer relative to the tube axis plays little to no role in determining the helical angle (Fig. S4).

Since it takes energy to maintain membrane curvature, the ability of EHD4^ΔN^ to generate membrane tubes smaller than its spontaneous curvature means that there is some additional factor(s) promoting curvature. An obvious candidate is the membrane interaction of EHD4^ΔN^. An alternative is the interaction of helices α1 and α12 from neighboring EHD4 monomers (interface-4), which mediates the packing of neighboring filaments into a continuous coat (Fig. 4C, D). Interface-4 shows variation as the radius of the tube varies, and preferences toward a particular orientation would influence the underlying tube radius.

## Discussion

Recent advances in cryo-EM have facilitated the structural analysis of membrane-bound protein scaffolds. Helical reconstructions requiring highly homogeneous samples (reviewed in (*38*)) have allowed, amongst others, medium to high resolution structural elucidation of the acetylcholine receptor (*39*), BAR domain proteins (*40, 41*), ESCRT (*42*), light-dependent protochlorophyllide oxidoreductase (LPOR) (*43*) and dynamin (*44*) assembled on membrane tubes. Structures of highly heterogeneous membrane-bound protein coats cannot be determined by helical reconstructions. However, recent advances in image processing have facilitated structure solution of such specimen by cryo-ET analysis combined with subtomogram averaging. Examples include structure determination of the membrane-bound COPI (*45*) and COPII coats (*46*), the N-BAR protein Bin1 (*47*) and the retromer (*48*). EHD4 samples bound to membrane tubes were highly heterogeneous (*32*), necessitating the use of cryo-ET for structural analysis. By projecting the obtained subtomogram averages back to the membrane tubes, this analysis allowed us not only to determine the structure of EHD4 within one filament but also to determine the EHD4 filament structures on various membrane curvatures. Our analysis has important implications for understanding how the ATPase cycle of EHDs is coupled to membrane recruitment, filament assembly and disassembly and how EHD4 generates membrane curvature. A resulting working model for the ATPase-dependent membrane cycle is outlined in the following.

Previous X-ray crystallographic analyses identified two conformations of EHDs. In the reported EHD2 crystal structure (*22*), the protein adopts a closed conformation whereas the crystal structure of EHD4 features an open conformation (*32*). Spectroscopic studies (*33*) suggest that EHDs are recruited to flat bilayers in an open conformation. Since the membrane interaction involves mostly polar interactions in the helical domain in the open conformation (*32*), EHDs may be in a rapid exchange with the cytosol. The G-domain is close towards the membrane in the open conformation so that the N-terminus can easily switch from its hydrophobic G-domain pocket into the membrane bilayer. The release of the N-terminus allows the KPF loop to enter the hydrophobic pocket to create oligomerization interface-2. Furthermore, our previous crystallographic study on EHD4 indicated that the G-interface (interface-3 in this manuscript) cannot be formed between EHD4 dimers in the open conformation due to steric constraints (*32*).

The transition of the open to the closed conformation, as observed in our study in the membrane-bound form, appears to be driven by the assembly of EHD oligomers onto curved membranes (Fig. 5A). By bilayer coupling (*49, 50*), the insertion of the hydrophobic helical tip region into the membrane is expected to generate positive membrane curvature. Similar to FYVE and ENTH domains (*51*), EHD membrane binding site is composed of charged residues and a hydrophobic membrane-penetrating protrusion. In turn, curved membranes may promote the transition from the open to the closed conformation in the EHD filament.

**Figure 5:**
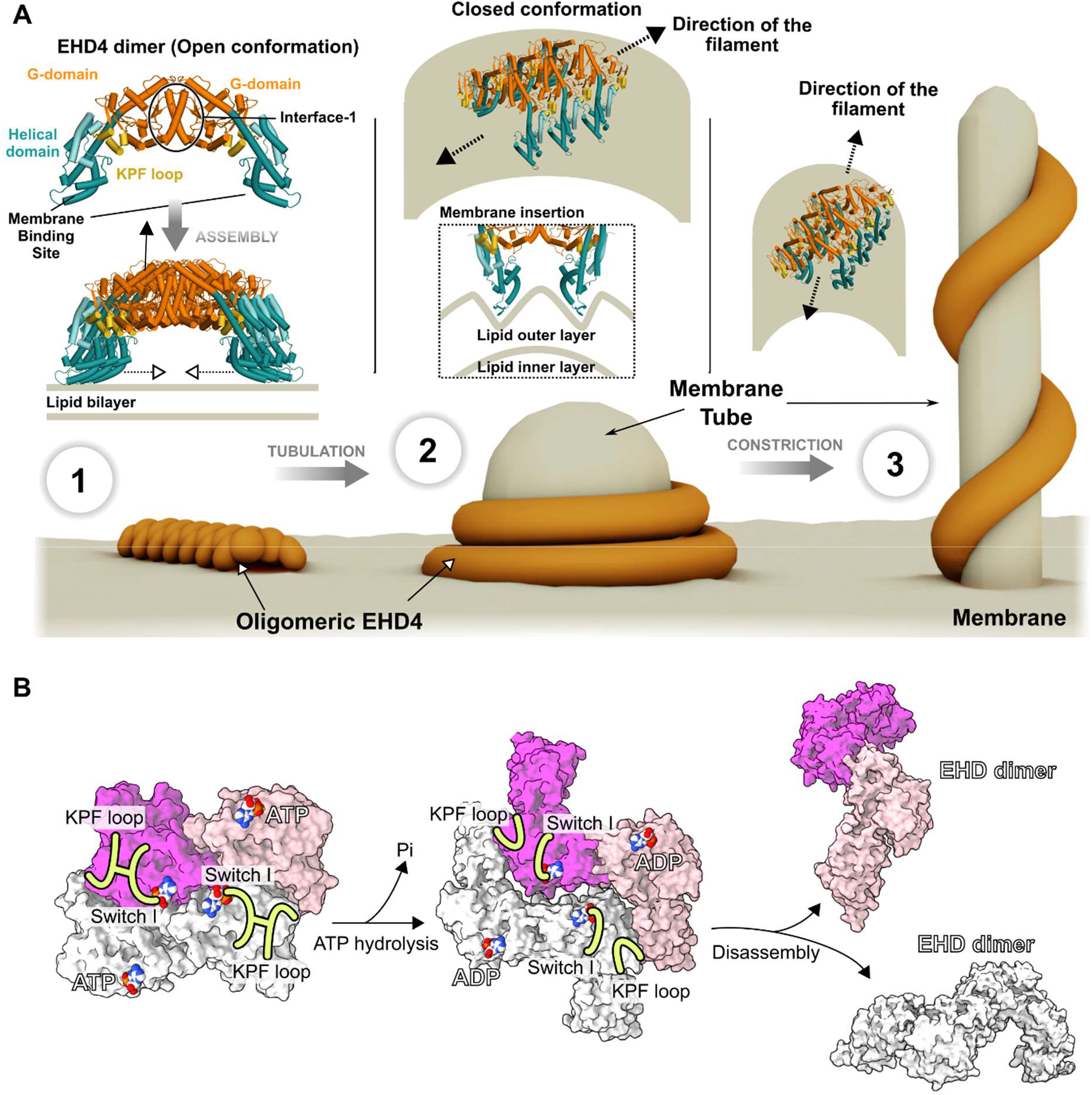
Model for EHD membrane remodeling: 1) ATP-bound EHD dimers are recruited to flat membranes in the open conformation where they oligomerize via interface-2 into filaments of low curvature. 2) Membrane curvature induces the transition of the open to closed conformation. In turn, insertion of the membrane binding site into the membrane promotes membrane curvature, which is associated with the formation of a stable helical filament via interface-3. See also Movie S3. 3) Constriction of the membrane tubule leads to increase of the helical pitch, which may allow interaction partners to be recruited to stabilize the membrane tubule. **B**. ATP hydrolysis leads to destabilization of the G interface and dissociation of the filament. The ADP-bound EHD dimer may convert back to the open conformation and dissociate from the membrane.

Upon initial curvature generation by wedging mechanism, EHD filaments then assemble into ring-like or helical oligomers via interfaces-2 and -3 (Fig. 5A), representing our membrane-bound structure. The involvement of the G-interface in this oligomer explains the strict ATP dependence of assembly (*23, 25*). ATP-binding stabilizes the switch regions which promotes dimerization (*34*). G-interface formation is associated with the displacement of the EH domain tail from the G-interface by the movement of the EH domains towards the periphery of the EHD filament (Fig. S3). In this orientation, the EH domains may bind to NPF-motif containing partner proteins (Fig. 5A and S3E, F). Formation of the G-interface is accompanied by a stimulation of the slow ATP hydrolysis reaction in EHDs (*21, 22*). In this way, ATP hydrolysis acts as an intrinsic timer to disassemble the EHD scaffold: In the ADP-bound state, the switch regions are destabilized and the interaction of switch I with the KPF loop is reduced (*32*) (Fig. 5B). Accordingly, interfaces-2 and -3 are weakened in the ADP-bound state, leading to dissociation of the oligomer (Fig. 5B). The ADP-bound EHD dimer may convert back to the open conformation and eventually dissociate from the membrane, therefore completing the ATPase cycle (Fig. 5B).

The EHD4 filament coat differs from the canonical oligomer architecture shared by several dynamin-related proteins, e.g. dynamin, DRP1, and Mgm1/OPA1, where stalk interactions define the interfaces forming the filament and nucleotide-dependent interactions between G-domains stabilize the inter-filament packing. EHDs on the other hand, orient the nucleotide-dependent interface along the filament direction and pack neighboring filaments with a stalk-stalk-like interaction (interface-4). This change in architecture allows the intrinsic filament curvature to be modulated by nucleotide, or perhaps more interesting, nucleotide state to be affected by the geometry of the filament. Additionally, whereas the pitch during active dynamin constriction is essentially fixed by the strong cross-filament interaction (*52*), EHD4 architecture may allow continuous deformation as the underlying membrane curvature changes.

As the helix angle of the filament changes, the orientation of the individual EHD4 dimers changes. This orientation has been previously suggested to be meaningful (*22*) as the membrane binding surface of EHD2 dimers in the closed conformation appears curved. Our structure shows that the dimer curvature is approximately 60° out of phase with the filament, meaning that when the helix angle is 60°, the dimer’s curvature is aligned with the tube curvature. In this orientation, any curvature generation by the dimer should be maximized. Interestingly, the maximum helical angles observed are roughly 60°, where the tube radius is about 15 nm. This state, with a large pitch, may represent the maximum curvature that the EHD4 filament can stabilize. In contrast to EHD4, ring-like assemblies were demonstrated for EHD1 (*25*) and EHD2 (*22*), likely indicating stronger spontaneous curvature for these oligomers.

Taken together, our structural analyses of the membrane-bound EHD4 scaffold elucidates novel insights into the coordination of the ATPase cycle with membrane recruitment, assembly and disassembly of the protein scaffold, and provides experimental insights into how membrane curvature is generated by EHD scaffolds.

## Methods

### Protein purification

Mouse EHD4 (residues 22-541, EHD4^ΔN^) and the indicated mutants were expressed from modified pET28 vector as N-terminal His6-tag fusions followed by a PreScission protease cleavage site. Expression plasmids were transformed in E. coli host strain BL21(DE3)-Rosetta2 (Novagen). Cells were grown at 37 °C in TB medium, and protein expression was induced at an optical density of 0.5 by the addition of 40 μM isopropyl-β-D-thiogalactopyranoside (IPTG), followed by overnight incubation at 18 °C. Upon centrifugation, cells were resuspended in resuspension buffer (50 mM Hepes/NaOH (pH 7.5), 500 mM NaCl, 25 mM imidazole, 2 mM MgCl_2,_ 2.5 mM β-mercaptoethanol (β-ME), 1 mM Pefabloc (Carl Roth), 1 µM DNase I (Roche)) and lysed in a microfluidizer. Following centrifugation (30,000 g, 1 h, 4 °C), cleared lysates were applied to a NiNTA column. The column was then extensively washed with washing buffer (50 mM Hepes/NaOH (pH 7.5), 700 mM NaCl, 10 mM CaCl_2_, 1 mM ATP, 10 mM MgCl_2,_ 10 mM KCl) and afterwards with equilibration buffer (50 mM Hepes/NaOH (pH 7.5), 500 mM NaCl, 25 mM imidazole, 2 mM MgCl_2,_ 2.5 mM β-ME). The protein was eluted with elution buffer I (50 mM Hepes/NaOH (pH 7.5), 500 mM NaCl, 2 mM MgCl_2,_ 2.5 mM β-ME, and 300 mM imidazole). Following the addition of 150 μg PreScission protease per 5 mg of protein, the protein was dialyzed overnight against dialysis buffer (50 mM Hepes/NaOH (pH 7.5), 500 mM NaCl, 1 mM MgCl_2_ and 2.5 mM β-ME). Following re-application of the protein to a NiNTA column to remove the His-tag, the protein was eluted with elution buffer II (50 mM Hepes/NaOH (pH 7.5), 500 mM NaCl, 2 mM MgCl_2_, 2.5 mM β-ME, and 50 mM imidazole). The uncleaved protein was concentrated using 30 kDa molecular weight cut-off concentrators (Amicon) and applied to a Superdex 200 gel filtration column equilibrated with SEC buffer (50 mM Hepes/NaOH (pH 7.5), 500 mM NaCl, 1 mM MgCl_2_, and 2.5 mM β-ME). Fractions containing the EHD4 constructs were pooled, concentrated and flash-frozen in liquid nitrogen. The purified protein was nucleotide-free, as judged by HPLC analysis.

### Liposome preparation

Liposomes were prepared by mixing 50 μL of Folch liposomes containing 50% Folch extract from bovine brain fraction I, 40% phosphatidylethanolamine and 10% cholesterol to 200 μL of a chloroform/methanol (1:0.3 v/v) mixture and dried under an argon stream. The liposomes were resuspended in liposome buffer (20 mM Hepes/NaOH (pH 7.5), 150 mM NaCl and 2.5 mM β-ME), sonicated in a water bath for 30 sec and extruded to 1 μm filter.

### Tubulation assay

For membrane tubulation assays, 10 μM EHD4^ΔN^ in tubulation buffer was incubated at room temperature for 20 min with 1 mg/ml liposomes.

### Cryo-electron microscopy and image processing

Complexes formed of EHD4^ΔN^ and liposome were diluted with a buffer containing 10 nm colloidal gold. 4 µl of this mixture was applied on a glow-discharged Quantifoil R2/2 grid (Quantifoil Micro Tools GmbH) and flash-frozen in liquid ethane using a Vitrobot Mark II device (FEI). The grids were stored under liquid nitrogen conditions until usage. Initial data was recorded on an in-house Thermo Fisher Scientific Talos L120C microscope operating at 120 kV on a Ceta Detector. The final set of 56 tilt series was imaged on a Thermo Fisher Scientific Titan Krios electron microscope operated at 300 kV equipped with a Gatan Quantum energy filter (slit with 20 eV) and a Gatan K2 detector using SerialEM (*53*). The nominal magnification was 53,000 x resulting in a pixel size of 2.628 Å. The data was acquired at a tilt range from -60 to 60 degrees using a dose-symmetric tilt scheme (*54*) at 3° increment. Tilt series were recorded as movies of 12 frames in counting mode and a dose rate of 2.3 e^-^/Å^2^ at defocus range of -3 um to -6 um resulting in a total dose of 94 e^-^/Å^2^ per tilt series. The initial contrast transfer function defocus value for each image of the tilt series was estimated using CTFFind4 (*55*). CTF correction was carried out by phase flipping using the program ‘ctf phase flip’ of the IMOD software package (*56*). Two copies of each tomogram were reconstructed using weighted back projection and SIRT in IMOD (*56*).

### Subtomogram averaging

The workflow described uses a combination of Dynamo and bespoke Matlab scripts. Initial particle picking was done using filament tracer by assigning the center of each tube. First round of alignments was performed using SIRT-filtered reconstructed tomograms binned 3 times (7.884 Å/pix). Particles for each tube were extracted using a box size of 128 pixels, randomized along its azimuth, averaged and low-pass filtered to 40 Å to generate the initial template for 3 rounds of coarse alignments. Oversampled particles converging onto the same coordinate were removed using Dynamo’s separation in tomogram parameter. Each tube was aligned individually and sub-boxed along the membrane to generate a section of the tube (Fig. 1B). Averaged sections were merged, aligned to the template and low-pass filtered to 40 Å. Multi-reference analysis (MRA) was used to eliminate bad particles from the dataset. Next, CTF corrected subtomograms were extracted and aligned to low-pass filtered references. Iterations were carried out starting from binned 3X data, using a low-pass filter of 20 Å, angular sampling of 12°, allowing shifts of 47 Å, and refinements were gradually improved by decreasing the binning factor, using less stringent low-pass filters and finer angular sampling. Final refinement steps were carried out on unbinned data extracted in 128 voxel boxes, using a low-pass filter set at 8 Å, angular sampling of 4°, and shift limits of 10 Å. A total of 23,813 subtomograms contributed to the final average. Mask-corrected resolution assessment was carried out within the RELION (*57*) post-processing framework using a soft-edged mask around the central EHD4 tetramer (Fig. S2), yielding a resolution of 7.6Å at the 0.143 FSC cut-off. Local resolution estimation and local filtering were applied using Phenix Local anisotropic sharpening and Phenix Local resolution map (*58*).

### Flexible fitting

The fitting procedure is summarized in Movie S2. An atomic model consistent with the cryo-EM map was generated using MDfit (*59*). MDfit uses the cryo-EM map as an umbrella potential to bias (i.e. deform) an underlying structure-based model (SBM) (*60*) in order to maximize the cross-correlation between the experimental density and the simulated electron density. An SBM is a molecular force field that is explicitly, albeit not rigidly, biased toward a certain native structure. The SBM for fitting was the EHD2 crystal structure (4CID) with the sequence homology modeled by Swiss-Model (*61*) to that of EHD4 (residues 22-535). The portion of the SBM for the KPF loop (residue 114-137), which is missing from the EHD2 structure, is based on the EHD4 crystal structure (5MVF). Building the SBM from the crystal structure ensured that the resulting model was maximally consistent with the crystal conformation. This entailed no significant changes in structure as the sequences are highly similar and included a missing loop in the crystal structure (residues 424-442). A preprocessing step was then necessary to move the EH domains within the dimer into a cis positioning because 4CID placed the EH domains *in trans*. This involved only reorientation of the 424-442 loop, no other residue positions were changed. We refer to this dimeric structure as EHD4-init. Since the EH domain is missing, the SBM for the EH domain is generated from the EHD2 crystal structure (residues 443-538). An SBM using EHD4-init as the input structure was then generated using SMOGv2.3beta (*60*) with the template “SBM_AA” meaning all non-hydrogen atoms were explicitly represented.

The density corresponding to the central two dimers within the cryo-EM map was chosen as the constraint for MDfit, since this region had the best resolution. Relaxation of the SBM under the influence of the cryo-EM map is performed by molecular dynamics (MD), and, thus, requires an initial condition. Two EHD4-init were rigid body fit into the map using the “Fit in Map” tool of Chimera. In order to compensate for the missing neighbors on either side of two dimers, the translational symmetry of the filament was exploited. Two additional copies of EHD4-init were added, positioned on either side, placed such that each dimer-dimer interface was identical. Technically, this was performed by 1) measuring the transformation X between the two central dimers in VMD, 2) duplicating the central dimers, and 3) applying X or -X to the duplicates. This four-dimer system served as the initial condition for MD. During MD, the duplicates were given strong position restraints, while the only constraint on the central dimers was the MDfit umbrella potential based on the cryo-EM map. Every 10^4^ MD steps, the duplicate dimers were repositioned. Through this iterative process, the structure converged within 3×10^5^ steps. The middle two dimers were taken as the atomic model. Note that even though the filament’s local C2 rotational symmetry was not explicitly enforced by us during MD, the fact that the SBM was based on a C2 symmetric structure ensured that this symmetry was included.

### Elastic model fitting

The fitted helical angle as a function of radius was defined by

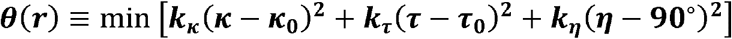

where the bit in brackets is an elastic energy and the min returns ***θ*** such that the radius is ***r*** and elastic energy is minimized. **κ** is the curvature, ***τ*** is the twist, ***η*** is the dimer orientation with respect to the tube axis, ***θ*** is the helix angle, and ***r*** is the radius of the tube. **κ**_0_ and ***τ***_0_ are the spontaneous curvature and twist. Fig. 4 shows the case for ***k***_***κ***_ **= 1, *k***_***τ***_ **= *k***_***η***_ **= 0**, where then twist elasticity and the dimer orientation are both negligible compared to filament curvature. Fig. S4 shows two alternatives, the first with ***k***_***κ***_ ***= k***_***τ***_**= 1, *k***_***η***_**= 0**, which is typical for continuous filaments, and the second with ***k***_***κ***_**= 1, *k***_***τ***_ **= 0, *k***_***η***_***=* 0.2**, *which* accounts for a preference for the dimer curvature to align perpendicular to the tube axis. For a constant helix, 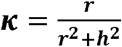, and 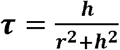, and 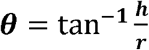, where *r* is the radius and 2π*h* is the pitch. In our EHD4 filament, the dimer is oriented approximately 30° relative to the helix angle, defining ***η*** = ***θ*** + **30** °. Therefore, when = **90**° = ***θ*** = **60**°, the dimer’s footprint curvature is optimally oriented with respect to the membrane tube. The two best fits with ***k***_***η***_ ***=0*** are performed using the Python library scipy.optimize.least_squares with, **κ**_**0**_ and ***τ***_**0**_ as fitting parameters. The curve with ***k***_***η***_**= 0. 2** uses, 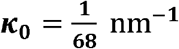 and has no free parameters to be fitted. Note that there is an analytic form for the minimum energy line with ***k***_***κ***_ ***=* 1, *k***_***τ***_ ***= k***_***η***_ ***=* 0**, which is given by 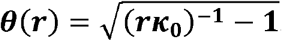.

### Code availability

Ad hoc scripts are available on Github: https://github.com/aamelo

## Acknowledgements

This work was supported by grants from the Deutsche Forschungsgemeinschaft (SFB 958/A12 to O.D. and Z3 to C.S.), the ERC grant MitoShape (ERC-2013-CoG-616024 to O.D.), a Humboldt fellowship to J.K.N. and the iNEXT grant PID3536 VID5570. We would like to thank Wim Hagen for support during cryo-ET data collection.

## Contributions

AM purified protein constructs, performed reconstitution of protein on membranes, optimized cryo-EM samples, processed data, determined the cryo-ET structure and analyzed the models. TS screened cryo-EM samples and preprocessed data. JN performed model fittings and analyses. JL assisted in data preprocessing. EVS and CH cloned, purified protein constructs, and performed membrane-binding assays. CS and OD supervised the project. AM, JN and OD wrote the article, with input from all authors.

## Competing interests

The authors declare no competing interests.

**Figure S1:**
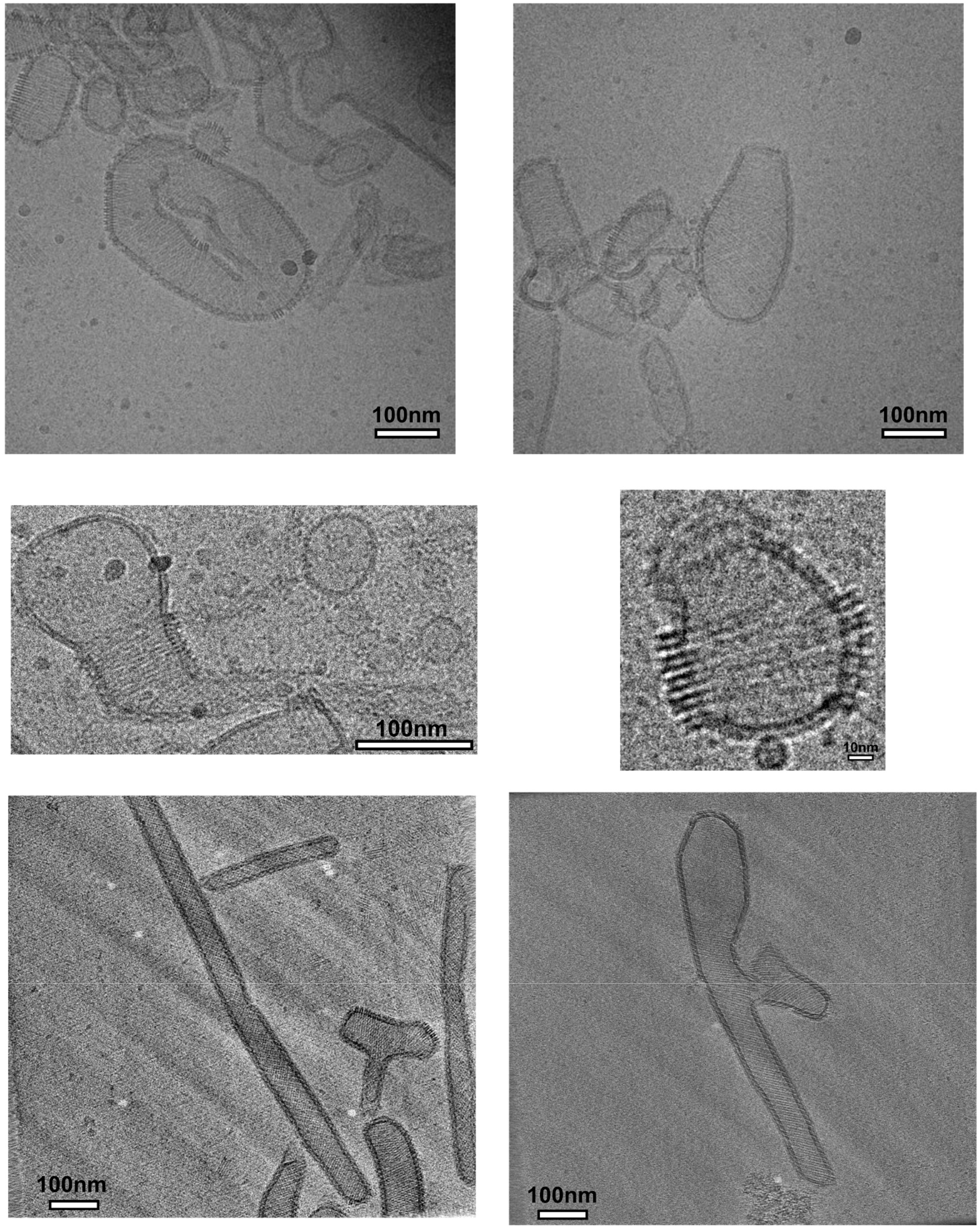
Heterogeneity of EHD4-coated membrane assemblies. Cryo-EM micrographs showing various examples of EHD4-coated membrane structures.

**Figure S2:**
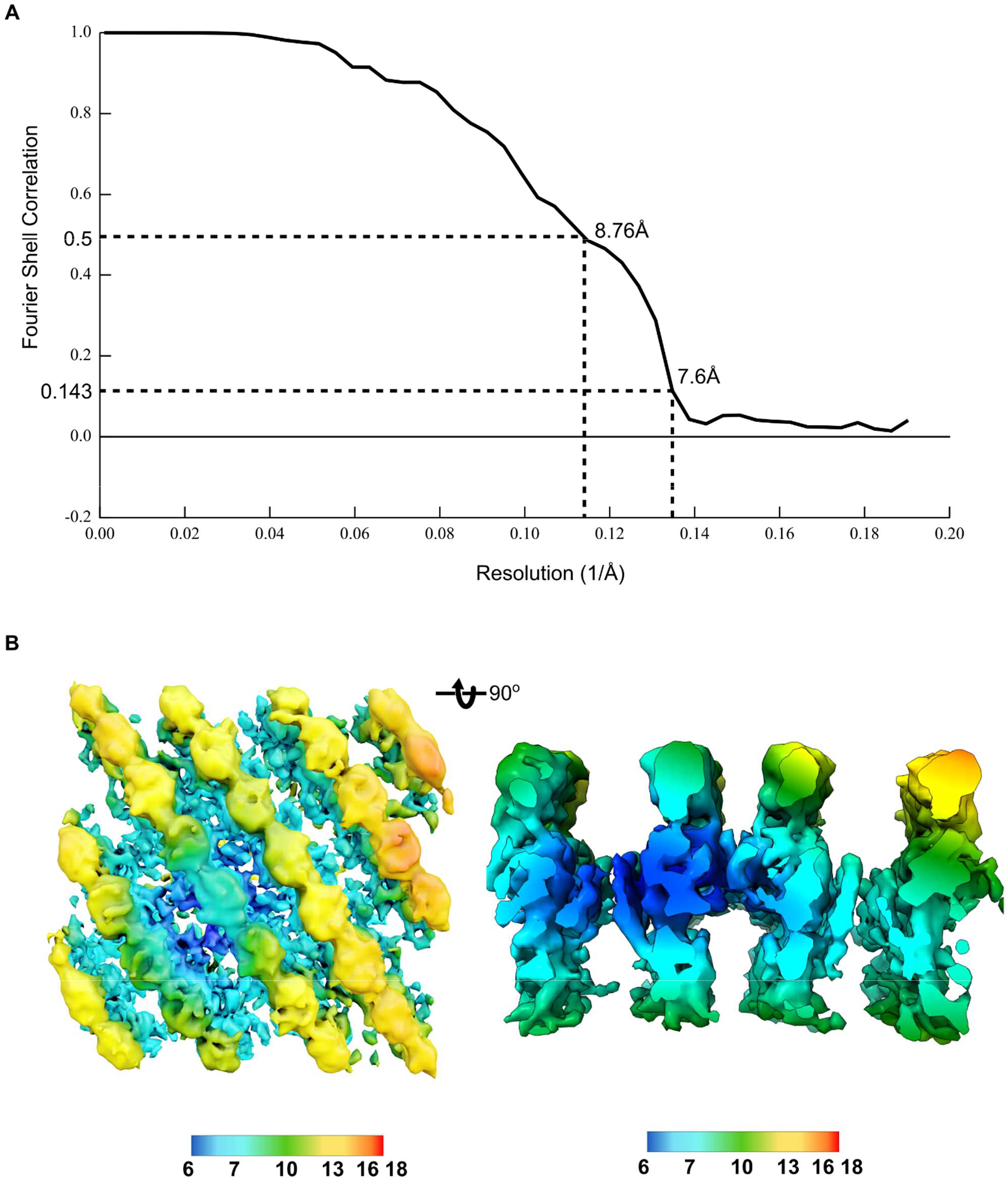
Resolution estimates. **A**. Mask-corrected gold standard Fourier Shell Correlation. **B**. Local resolution estimated with Phenix (*58*).

**Figure S3.**
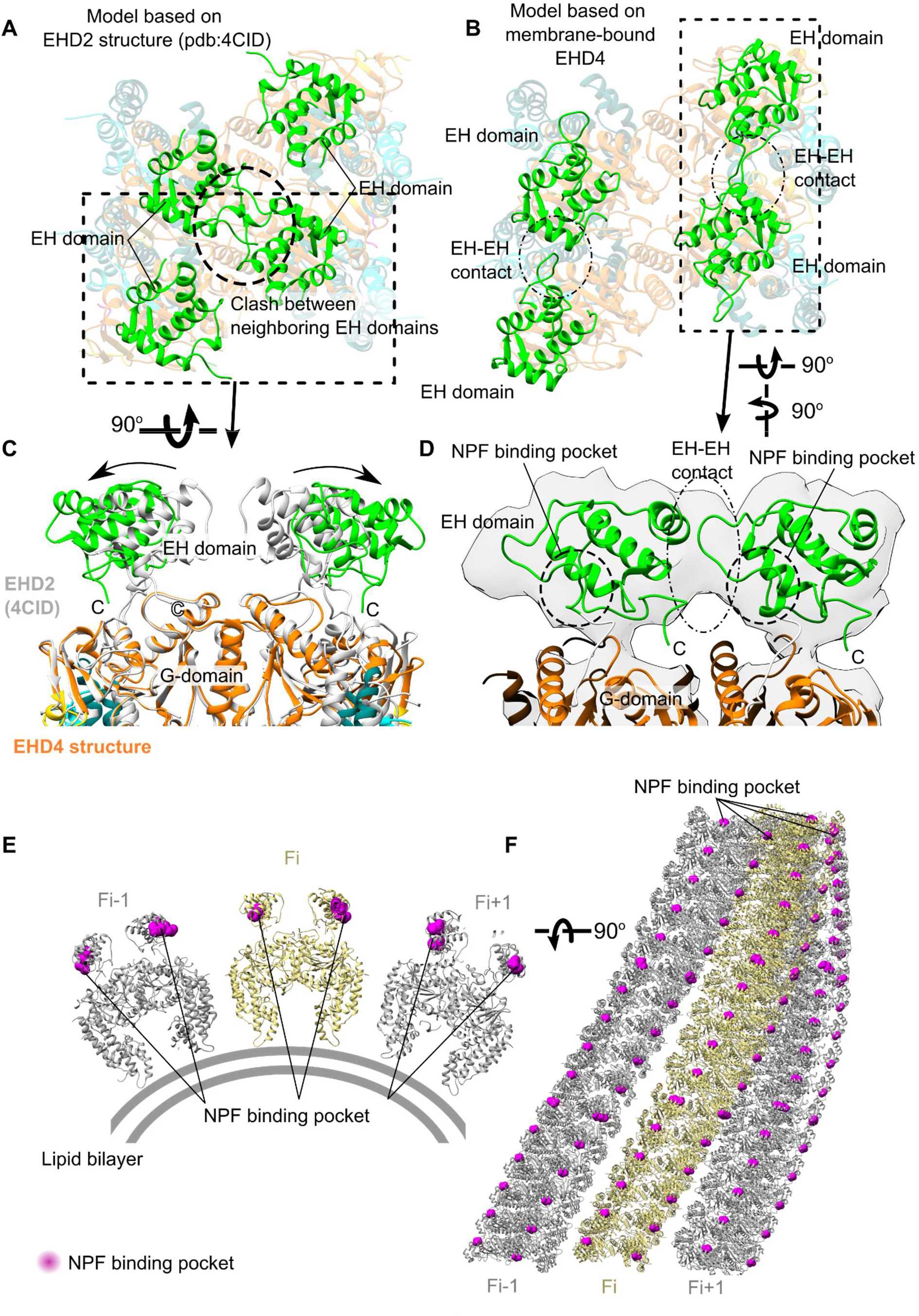
Displacement of the EH domains. **A**. The position of the EH domain was modelled in the filament, based on the EHD2 crystal structure. In this position, EH domains of adjacent dimers would clash. **B**. EH domains of adjacent dimers contact each other in the membrane-bound EHD4 structure. **C**. Superposition of the membrane-bound EHD4 and EHD2 crystal structure dimer (grey). Upon membrane binding, the EH domains (green) move towards the periphery of the filament. The C-terminus of the EH domains folds into the nucleotide-binding pocket of the G-domains in the EHD2 crystal structure but is displaced in the membrane-bound structure. **D**. Side view on the filament showing the new EH domain contact. Frontview (**E**) and topview (**F**) of NPF binding pockets highlighted in the EHD4 filament. They are positioned on the outer region of the EHD4 filament and are available for interactions with NPF-containing proteins at the cytosol or at the membrane.

**Figure S4.**
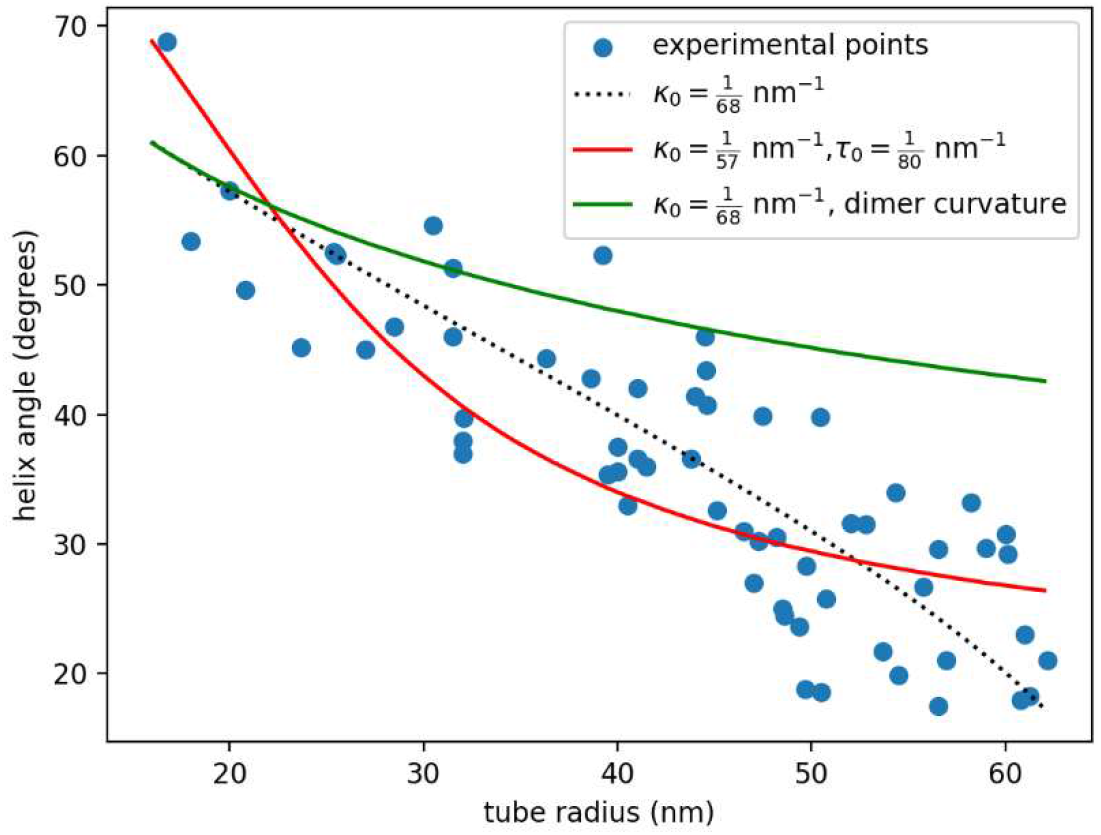
Two additional models compared to experimental points. Dotted line shows best fit result of main text Fig. 4. Red line shows best fit when taking the curvature and twist moduli as equal. While the fit is not obviously wrong, the better fit of the dotted line (where twist modulus is zero) suggests the twist modulus is relatively small. The green line shows the curve given by the dotted line plus an addition accounting for an alignment driven by the dimer curvature. The large tube behavior significantly deviates. See Methods for details.

**Table S1:** Cryo-EM data collection, refinement and validation statistics.

| **DATA COLLECTION AND PROCESSING** |  |
| --- | --- |
| MAGNIFICATION | **53,000** |
| VOLTAGE (kV) | **300** |
| ELECTRON EXPOSURE (e/Å<sup>2</sup>) | **94** |
| DEFOCUS RANGE (μm) | **3-6** |
| PIXEL SIZE (Å) | **2.628** |
| SYMMETRY IMPOSED | **C2** |
| INITIAL PARTICLE IMAGES (NO.) | **84,000** |
| FINAL PARTICLE IMAGES (NO.) | **23,813** |
| MAP RESOLUTION (Å) | **7.6** |
| FSC THRESHOLD | **0.143** |
| MAP RESOLUTION RANGE (Å) | **6.47-16** |

**Movie S1.**
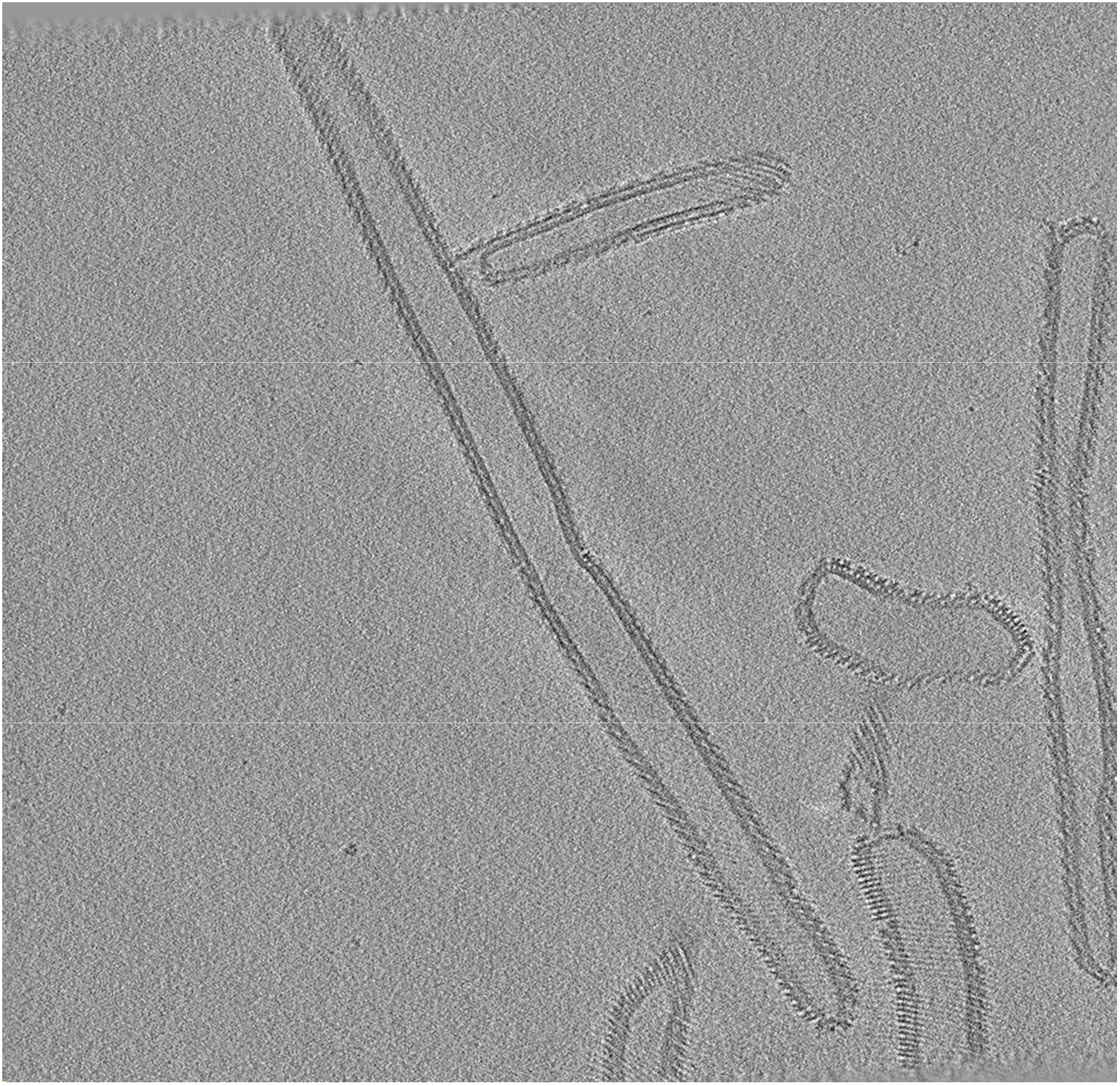
Orthogonal views of the reconstructed tomogram containing EHD4 coated tubes.

**Movie S2:**
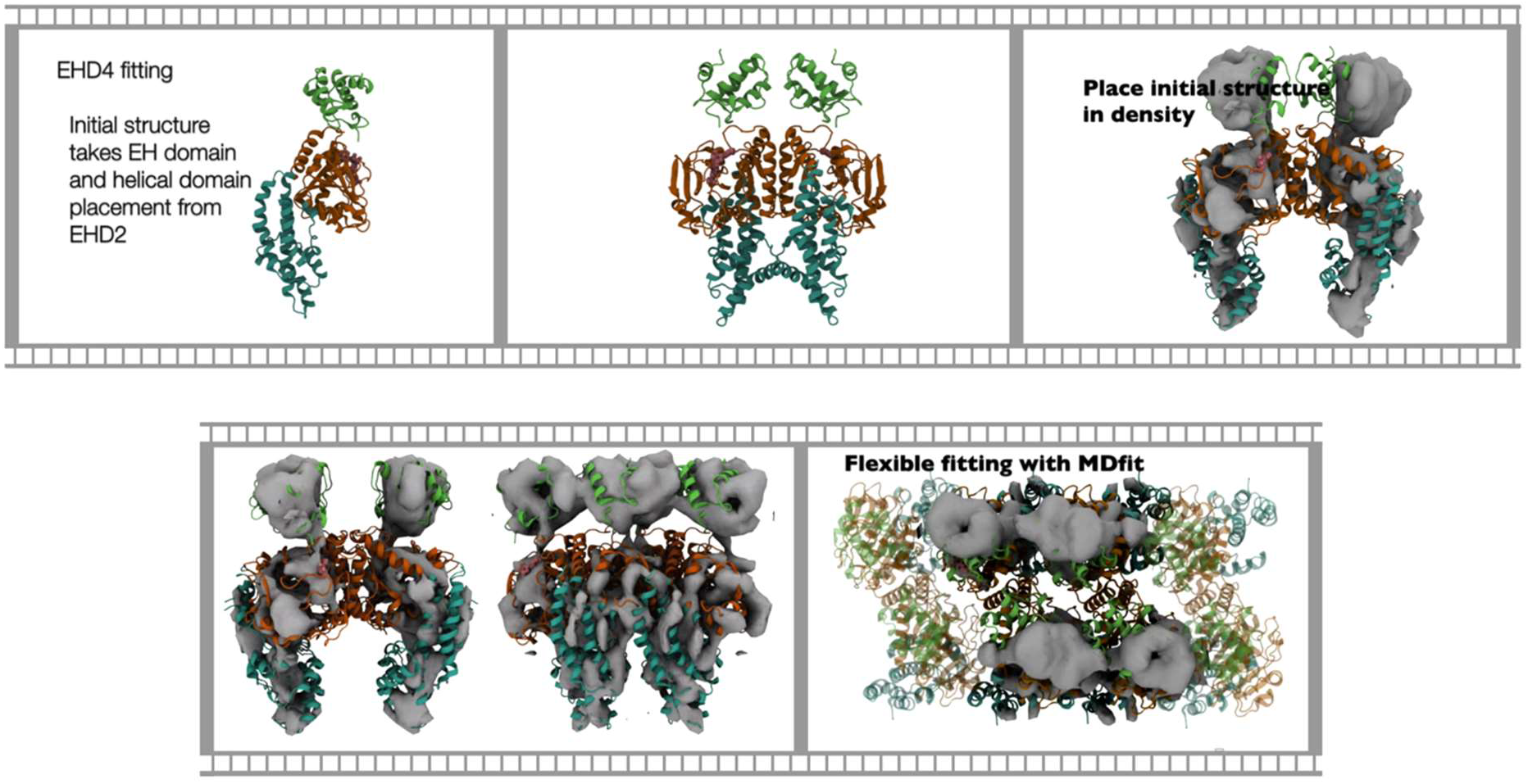
Flexible fitting of membrane-bound EHD4 based on the crystal structures.

**Movie S3.**
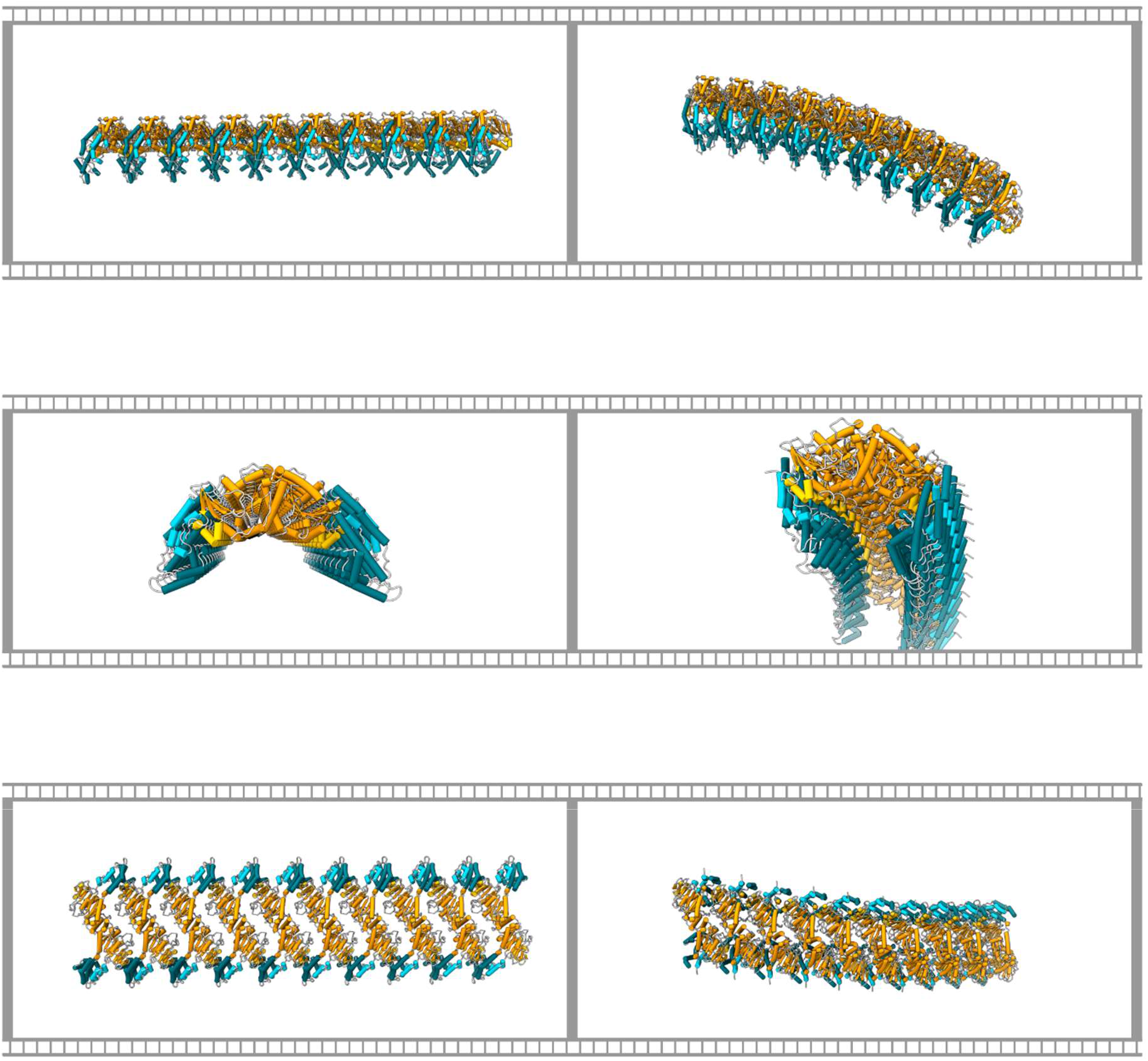
Transition between the open linear EHD4 filament to the curved closed filament.

